# Assessing the brain representation of conscious and unconscious visual contents using encoding based representational similarity analysis

**DOI:** 10.1101/2022.12.23.521727

**Authors:** Ning Mei, Roberto Santana, David Soto

## Abstract

The development of novel frameworks to understand the properties of unconscious representations and how they differ from the conscious counterparts may be critical to make progress in the neuroscience of vision consciousness. Here we re-analysed data from a within-subject, high-precision, highly-sampled fMRI study (N=7) coupled with model-based representational similarity analysis (RSA) in order to provide an information-based approach to study the representation of conscious and unconscious visual contents The standard whole-brain searchlight RSA revealed that the hidden representations of convolutional neural network models explained brain activity patterns in response to unconscious contents in the ventral visual pathway in the majority of the observers, particularly for models that ranked high in explaining the variance of the visual cortex (i.e., VGGNet and ResNet50). Also five of seven subjects showed brain activity patterns that correlated with the model in frontoparietal areas in the unconscious trials. However, the results of an encoding-based RSA analyses in the unconscious condition were mixed and somehow difficult to interpret, including negative correlations between the representations of the computer vision models and the brain activity in frontal areas in a substantial amount of the observers.

## Introduction

The main objective of this work is to provide an information-based approach to pinpoint the representation of unconsciously processed stimuli in human vision. Unconsciously processed stimuli are thought to be encoded transiently and locally in visual cortex ^1^. However, recent work has providing intriguing data suggesting that unconscious information processing may be implicated higher-order processes, namely, cognitive control ^2–4^, memory-guided behaviour ^5–9^, and even language ^10–12^. Follow-up work however did not find support for this view ^13^, and purported unconscious processing effects have been difficult to replicate ^14^.

The controversy on the study of unconscious processing is likely due the lack of sound frameworks to isolate it, both behaviourally and at a neural level ^15^. A recent study presented the first high-precision, highly-sampled, within-subject approach to pinpoint the neural representation of unconscious contents ^16^, by leveraging the power of machine learning and biologically plausible computational models of visual processing. Mei and colleagues ^16^ showed that unconscious contents, even those associated with null sensitivity, can be reliably decoded from multivoxel patterns that are highly distributed along the ventral visual pathway and also involving parieto-frontal substrates.

Here we used model-based representational similarity analysis (RSA) to provide a more fine-grained information-based approach to the study of unconscious and conscious neural representations in human vision. RSA quantifies the geometry of the multi-voxel representations using a collection of pairwise item dissimilarity measurements among BOLD responses in a given region of interest ^17,18^. RSA characterizes the information patterns that come from different aspects ^18^. The activity-pattern information could be measured in the so-called representational similarity analyses by distance-based measurements such as Euclidean distance ^19^. The distance-based approach compares the distance between pairs of activity patterns relative to a computational model. The distances indicate the relative similarities among pairs of activity patterns, including computational model patterns. Thereby allowing comparison across different levels of analyses and different principles of information processing.

The deep convolutional neural network is one of the classes of deep neural network models in computer vision that had been used to explain brain activity ^20–23^. In order to quantify the brain responses to the masked images, we compared the information patterns captured by the state-of-the-art computer vision models ^24^ and the information patterns captured by the fMRI. Thus, we conducted a standard RSA and an encoding-based RSA konkle2022self, which are explained below.

Observers (N = 7) performed six fMRI sessions across 6 days leading to a total of 1728 trials per subject. Observers were presented with gray-scaled images of animate and inanimate items with a random-phase noise background ^25^. The target images were presented briefly, preceded and followed by a dynamic mask composed of several frames of Gaussian noise. On each trial of the fMRI experiment, participants were required to discriminate the image category and to indicate their subjective awareness (i.e. (i) no experience/just guessing (ii) brief glimpse (iii) clear experience with a confident response). The duration of the images was based on an adaptive staircase that was running throughout the experiment (see Methods), and which, based on pilot testing, was devised to obtain a high proportion of unconscious trials. On average, target duration was (i) 25 ms on trials rated as unaware (ii) 38 ms on glimpse trials and (iii) 47 ms on the aware trials.

Figure 1 illustrates an example of a trial in the behavioural task. The signal detection theoretic analyses of the behavioural data can be found in Mei et al. (2022) ^16^. Basically, participants 1-4 showed null perceptual sensitivity of the stimulus category on the trials rated as unaware, while the participants 5-7 showed above chance discrimination performance on unaware trials.

**Figure 1:**
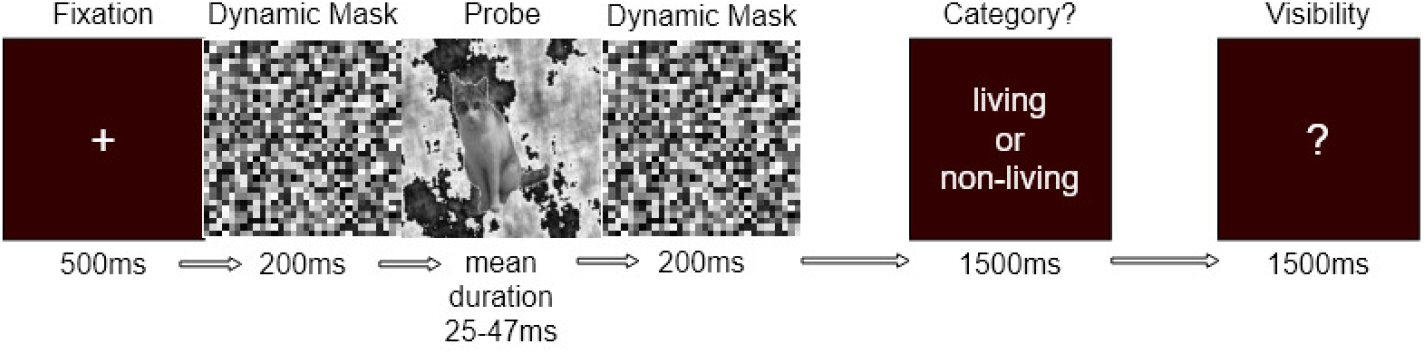
Example of the sequence of events within an experimental trial. Observers were asked to discriminate the category of the masked image (living vs. non-living) and then rate their visual awareness on a trial by trial basis. Example of a trial with a masked image of a cat.

## Methods

### Representations of the images from the state-of-the-art computer vision models

The information patterns captured by the computer vision models were quantified by the representational dissimilarity matrix (RDM, namely model RDM). First, we fine-tuned the pretrained models with the Caltech101 dataset ^26^ to control for the size of the hidden representations (i.e., to match the number of units across models for the RSA). A 2D adaptive average pooling was applied to the last convolutional layer of the FCNNs so that the outputs became one-dimensional vectors ^27^ ^*a*^. The pooled activity patterns of the Alexnet and the VGG19 had 512 units, MobilenetV2 and DenseNet169 had 1024 units, and ResNet50 had 2048 units. Then, a fully-connected layer with 300 units was added to the outputs of adaptive pooling, followed by a scaled exponential linear unit function (SELU, ^28^) ^*b*^. This allowed us to rescale the representational dimensions of different models to the same dimension. And SELU was chosen to speed up the training of the new components and scale the output activity patterns for better propagating information to the next layer.

A fully-connected layer with 96 units (1 per category of the Caltech dataset) was added to the outputs of the SELU layer, followed by a softmax activation function to compute probabilistic predictions of the image categories. The model was trained using 96 unique categories of the Caltech101 images (BACKGROUND_Google, Faces, Faces_easy, stop_sign, and yin_yang were excluded, ^26^). The convolutional layers were frozen during the training. The loss function was binary cross entropy and the optimizer was stochastic gradient descent. The data was split into train and validation partitions and the training was terminated if the performance on the validation data did not improve for five consecutive epochs.

We then fed the trained FCNNs with exactly the same images used in our experiment but without the noise background, in order to extract the hidden representations of the images. We then average the hidden representations of the trials belonging to the same item (i.e cat) and computed the model RDMs for the AlexNet, the VGGNet, MobileNetV2, the ResNet50, and the DenseNet169 (Figures 2,3, 4, 5, and 6 depict the RDMs for these hidden representations), and it appears to be clear clusters for the animate and inanimate image categories. Although it might be difficult to visually inspect for AlexNet, the animate images were highly similar among each other and the inanimate images were highly similar among each other, forming clusters at the lower left and upper right quadrants. The similarities were more consistent among animate images (lower left quadrant) compared to the inanimate images (upper right quadrant).

**Figure 2:**
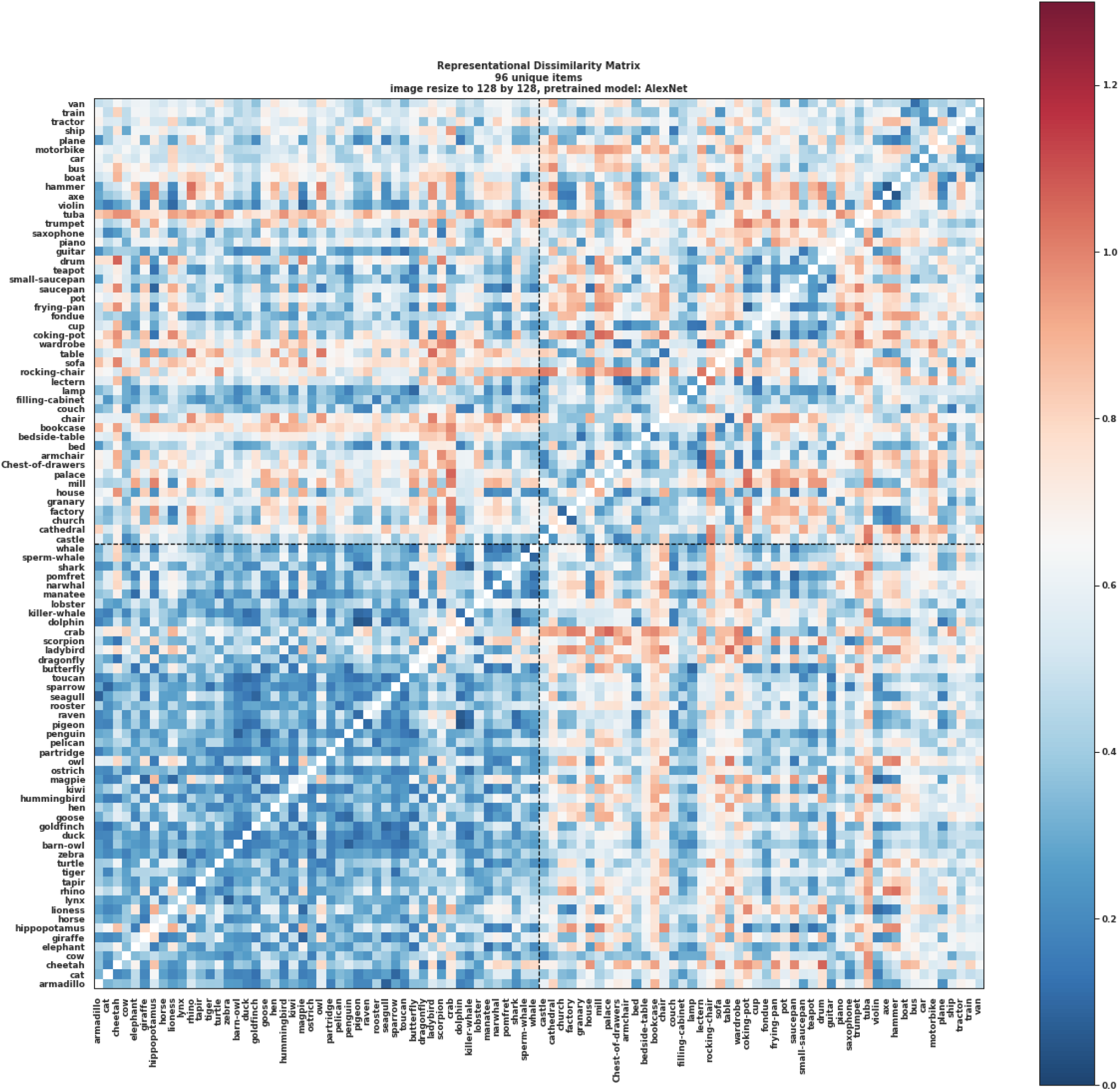
Representational dissimilarity matrix of the hidden representations of the AlexNet model fine-tuned by Caltech101 dataset. The images were resized to 128 × 128 × 3 and then passed through the model to obtain the feature representations. The RDM was computed by 1 - Pearson correlations of the feature representations. The first 48 items were animate and the last 48 items were inanimate.

**Figure 3:**
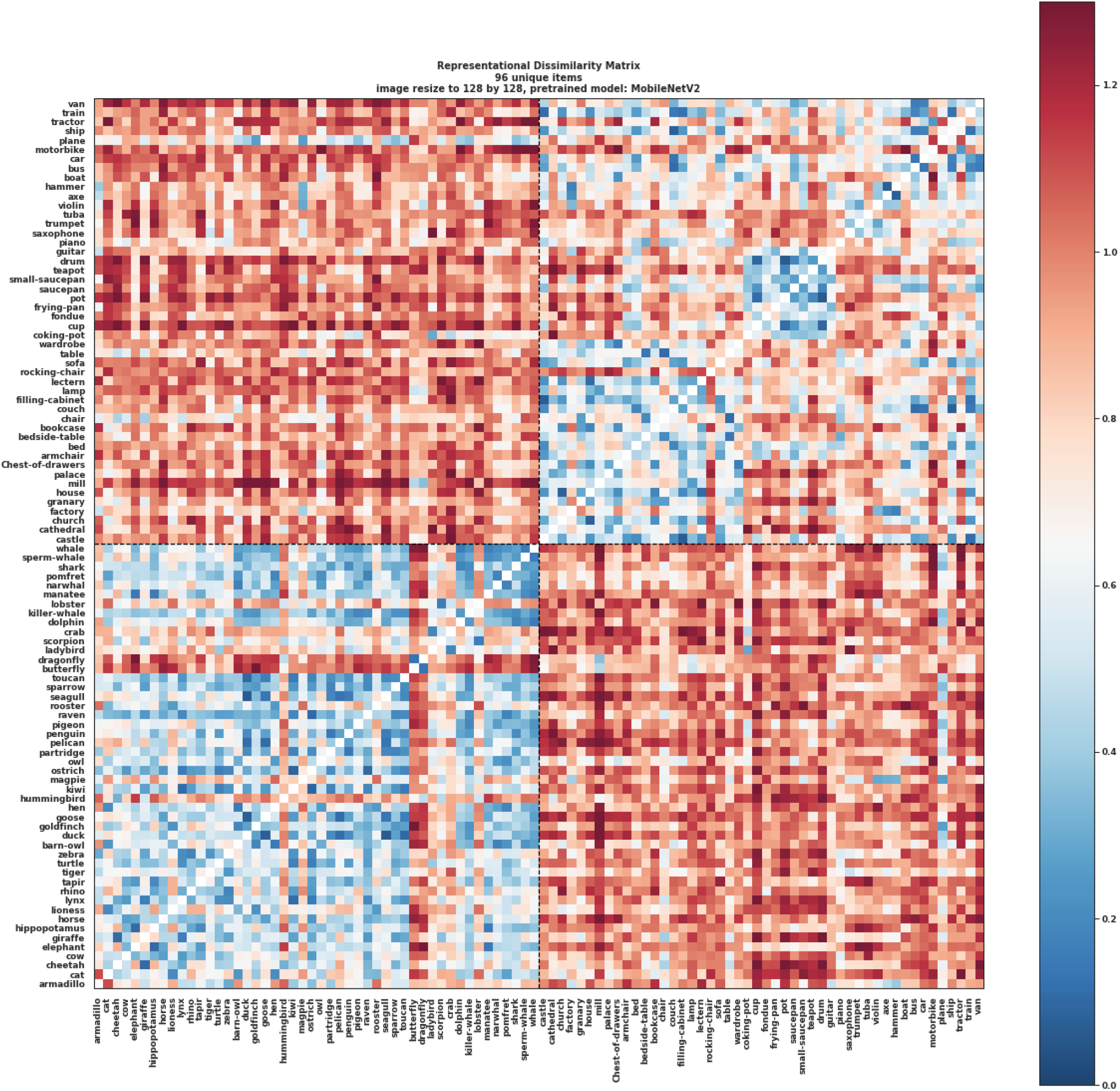
Representational dissimilarity matrix of the hidden representations of the MobileNetV2 model fine-tuned by Caltech101 dataset. The images were resized to 128 × 128 × 3 and then passed through the model to obtain the feature representations. The RDM was computed by 1 - Pearson correlations of the feature representations. The first 48 items were animate and the last 48 items were inanimate.

**Figure 4:**
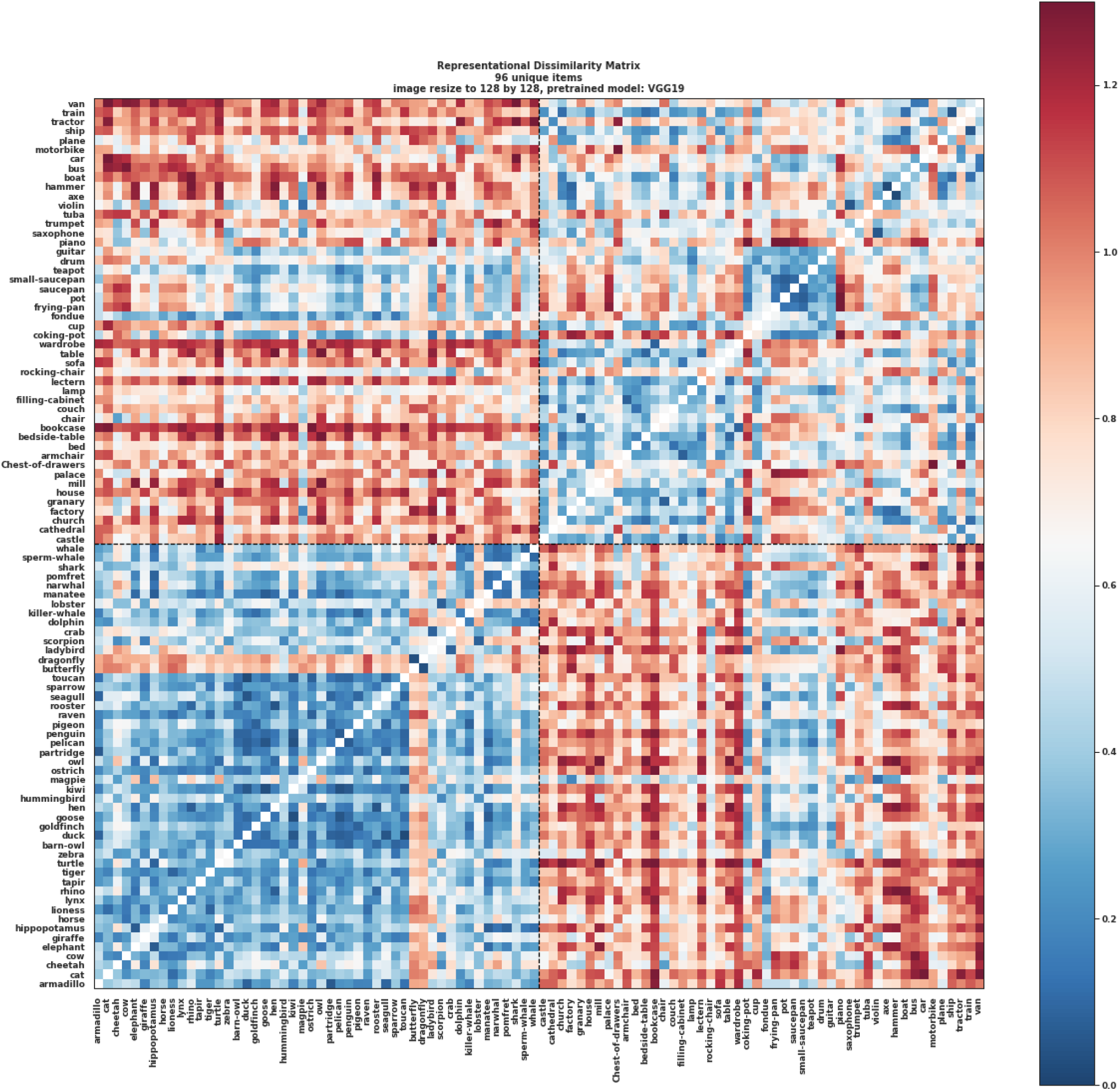
Representational dissimilarity matrix of the hidden representations of the VGG19 model fine-tuned by Caltech101 dataset. The images were resized to 128 × 128 × 3 and then passed through the model to obtain the feature representations. The RDM was computed by 1 - Pearson correlations of the feature representations. The first 48 items were animate and the last 48 items were inanimate.

**Figure 5:**
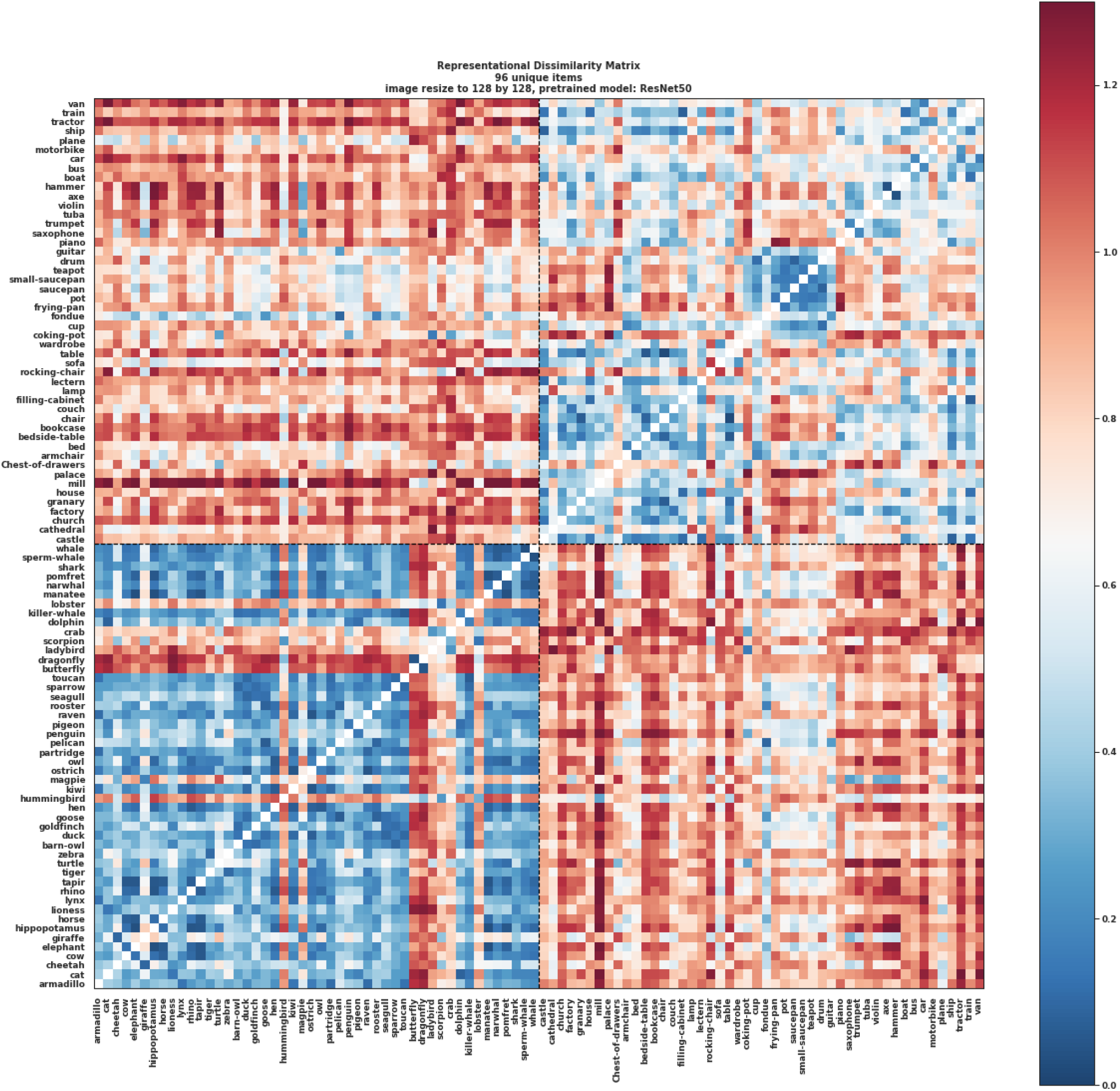
Representational dissimilarity matrix of the hidden representations of the ResNet50 model fine-tuned by Caltech101 dataset. The images were resized to 128× 128× 3 and then passed through the model to obtain the feature representations. The RDM was computed by 1 - Pearson correlations of the feature representations. The first 48 items were animate and the last 48 items were inanimate.

**Figure 6:**
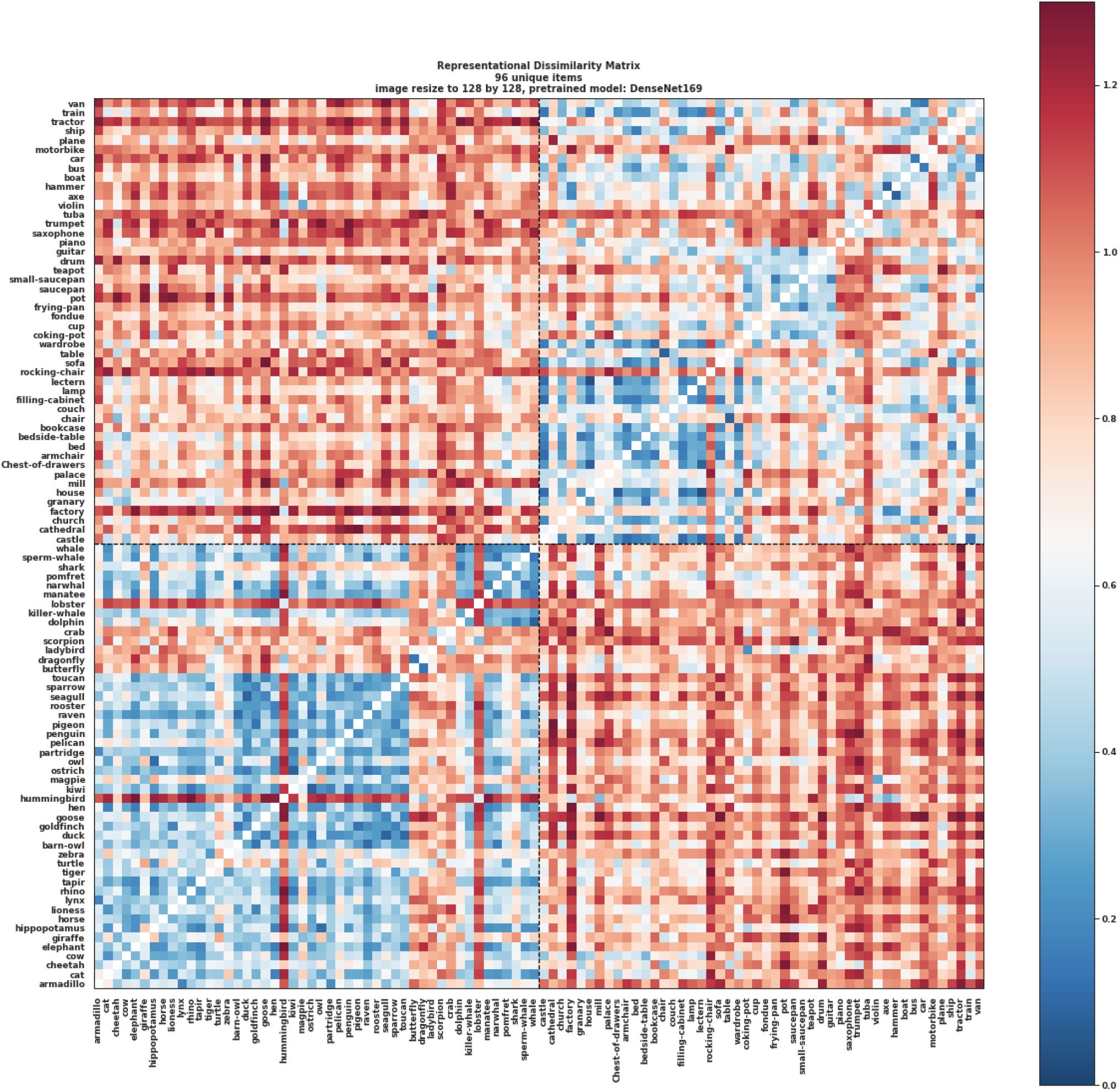
Representational dissimilarity matrix of the hidden representations of the DenseNet169 model fine-tuned by Caltech101 dataset. The images were resized to 128 ×128 ×3 and then passed through the model to obtain the feature representations. The RDM was computed by 1 - Pearson correlations of the feature representations. The first 48 items were animate and the last 48 items were inanimate.

Using these RDMs, we further conducted a standard RSA and an encoding-based RSA on the fMRI data within each subject.

**Figure 7:**
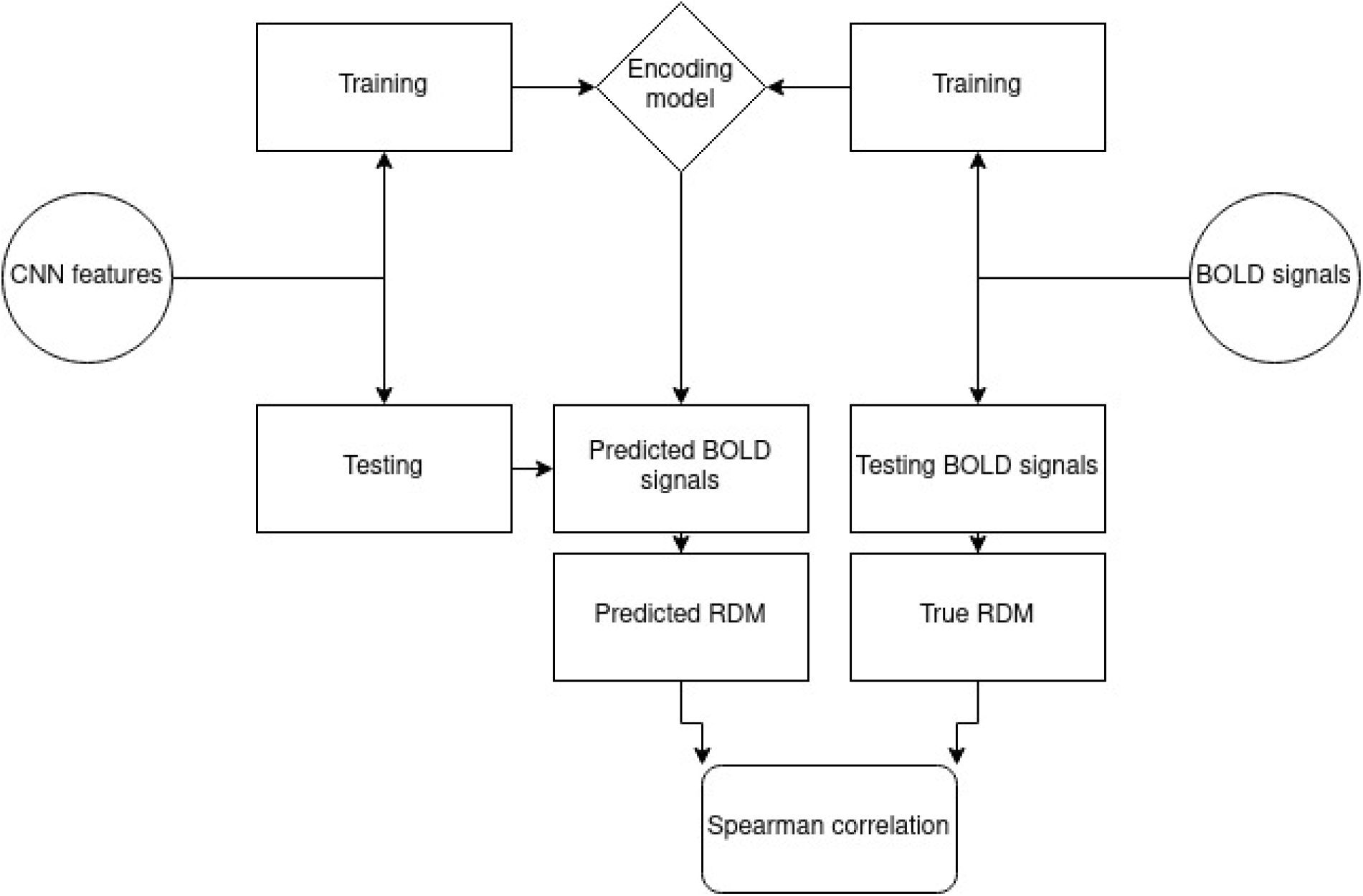
General diagram of the encoding-based RSA pipeline. The CNN features and the BOLD signals were split into “training” and “testing” sets accordingly for within- and cross-conscious state conditions. The training set was used to train the encoding model (Ridge regression), and the encoding model was used to predict the BOLD signals in the testing set using the testing set CNN features. In the end, we correlated the predicted BOLD signals with the true BOLD signals to obtain the Spearman correlation brain map for further analysis.

### Standard whole-bk

To conduct a whole-brain searchlight RSA, we then extracted the BOLD activity patterns using a mask that contained all the ROIs used for the decoding analyses. We averaged the BOLD signals of trials belonging to the same item (i.e., cat) within a searchlight sphere of 9mm that moved around the brain mask. An RDM was computed for the extracted BOLD signals using 1 - Pearson correlation implemented by Scipy ^29^ ^*c*^ among each pair of images within each sphere. The lower triangles of the two RDMs were extracted and correlated using Spearman correlation for each sphere of the searchlight. The Spearman correlation coefficient was assigned to the center of each sphere of the whole-brain searchlight. The resulting brain map of FCNNs/brain similarities was converted to standard space for visualization using Nipype ^30^ and FSL ^31^ tools. This pipeline was run separately in the unconscious and conscious conditions, separately within each participant.

### Encoding-based whole-brain searchlight RSA

Konkle and colleagues ^32^ proposed an encoding-based RSA pipeline, adding encoding models on top of the standard RSA pipeline, which can be used to better contextualize the information patterns from the computational model space (i.e., VGG19) to the brain space.

To conduct an encoding-based RSA, we split the data into train and test sets by leaving four unique items out. With 96 unique items, we had twenty-four folds for cross-validation. Within the training set, an L2-regularized linear regression model (ridge regression) implemented by scikit-learn ^33,34^ was trained to predict the voxel values using the image features in the computational model space. The image features in the computational model space were the information patterns captured by the computer vision models fine-tuned by the Caltech101^26^. The fine-tuning procedure is described in section. Prior to training the ridge regression model, two scalers were trained using only the training set. One scaler was a MinMaxScaler ^*d*^ that normalized the voxel values, and the other was a StandardScaler ^*e*^ that normalized the image features. These scalers were trained in the training set and then applied to the test set data for data normalization to improve the analyzing performance and robustness. The ridge regression model was nested in a grid-search algorithm to cross-validate the best L2-regularization term by stratified random shuffle splitting in the training set using twenty folds. The trained ridge regression model predicted the voxel values in the test set. The predicted voxel values were averaged for each unique item. An RDM of the predicted voxel values and an RDM of the original voxel values in the test set were computed using 1 - Pearson correlation implemented by Scipy ^29^ among each pair of items within each sphere. The lower triangles of the two RDMs were extracted and correlated using Spearman correlation for each sphere of the searchlight and the correlation coefficients were assigned to the center of the spheres.

During cross-validation, trials corresponding to one living (i.e., cat) and one non-living (i.e., boat) item for a given awareness state (i.e., unconscious) were left out as the test set and the rest was used to fit the machine learning pipeline. With 96 unique items, 2256 cross-validation folds could be performed in principle. However, because the awareness states were randomly sampled for each unique item (i.e., cat), the proportion of examples for training and testing were not equal among different folds. Some subjects had less than 96 unique items for one or more than one of the awareness states. Thus, less than 2256 folds of cross-validations were performed in these cases. The number of total available instances ranged from 700 to 800 and the number of test sets ranged from 2 (one trial for living and one trial for nonliving) to 20 for either unconscious or conscious states.

For cross-state encoding-based RSA, the encoder was trained from data in a particular awareness state (the “source”; e.g., on conscious trials) and then tested on a different awareness state (the “target”; e.g. on unconscious trials) on top of the cross-validation procedure described above. Similar to the within-state encoding-based RSA, instances corresponding to one living and one non-living item in both “source” and “target” were left out, but only the left-out instances in the “target” were used as the test set. The rest of the instances in “source” were used as the training set to fit the machine learning pipeline (preprocessing + encoder) as described above. Because the awareness states were randomly sampled for each unique item (i.e., cat), the proportion of examples for training and testing were not equal among different folds. The number of total available instances ranged from 700 to 800 and the number of test sets ranged from 2 (one trial for living and one trial for nonliving) to 20.

## Results

### Standard searchlight RSA results

In the unconscious condition (Figure 8), an RSA based on the AlexNet to extract the model representations of the images showed that six of seven subjects demonstrated model-relevant representations in both the ventral visual pathway and frontoparietal areas. An RSA based on VGG19 representations showed that four subjects had model-relevant representations in the ventral visual pathway, and a different subset of four subjects showed model-relevant representations in the frontoparietal areas. Using the MobileNet, five subjects showed model-relevant representations in the ventral visual pathway, and four subjects showed model-relevant representations in the frontoparietal areas. ResNet50-based RSA showed model-relevant representations in the ventral visual pathway in five subjects, and six subjects showed model-relevant representations in the frontoparietal areas. Finally, the DenseNet169-based RSA showed that three subjects had model-relevant representations in the ventral visual pathway, and four subjects showed model-relevant representations in the frontoparietal areas.

**Figure 8:**
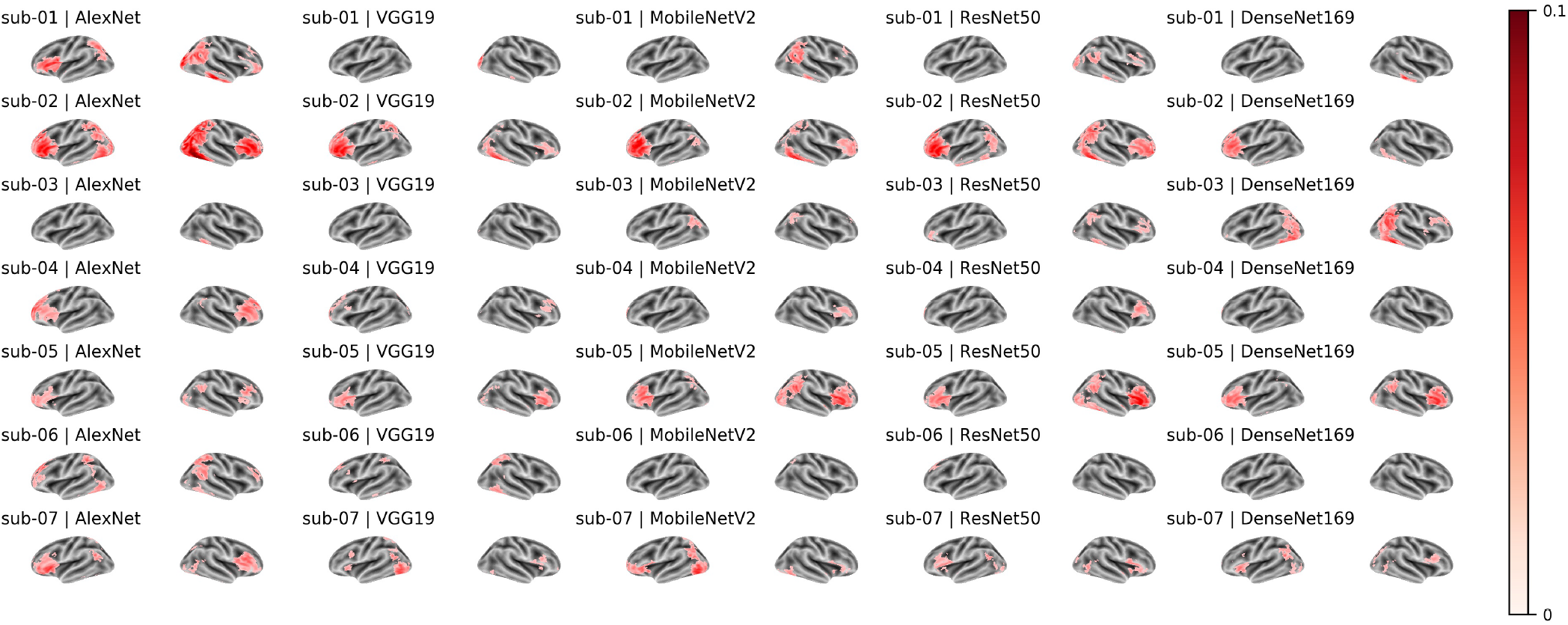
RSA map of the unconscious condition for the five FCNNs. Each row showed the left and right hemisphere RSA maps of the same subject, and every two columns showed the left and right hemisphere RSA maps of the same computer vision model.

For the conscious condition (Figure 9), all subjects showed model-relevant representations in the ventral visual pathway across all models. Six of seven subjects also showed model-relevant representations in the frontoparietal areas across all models.

**Figure 9:**
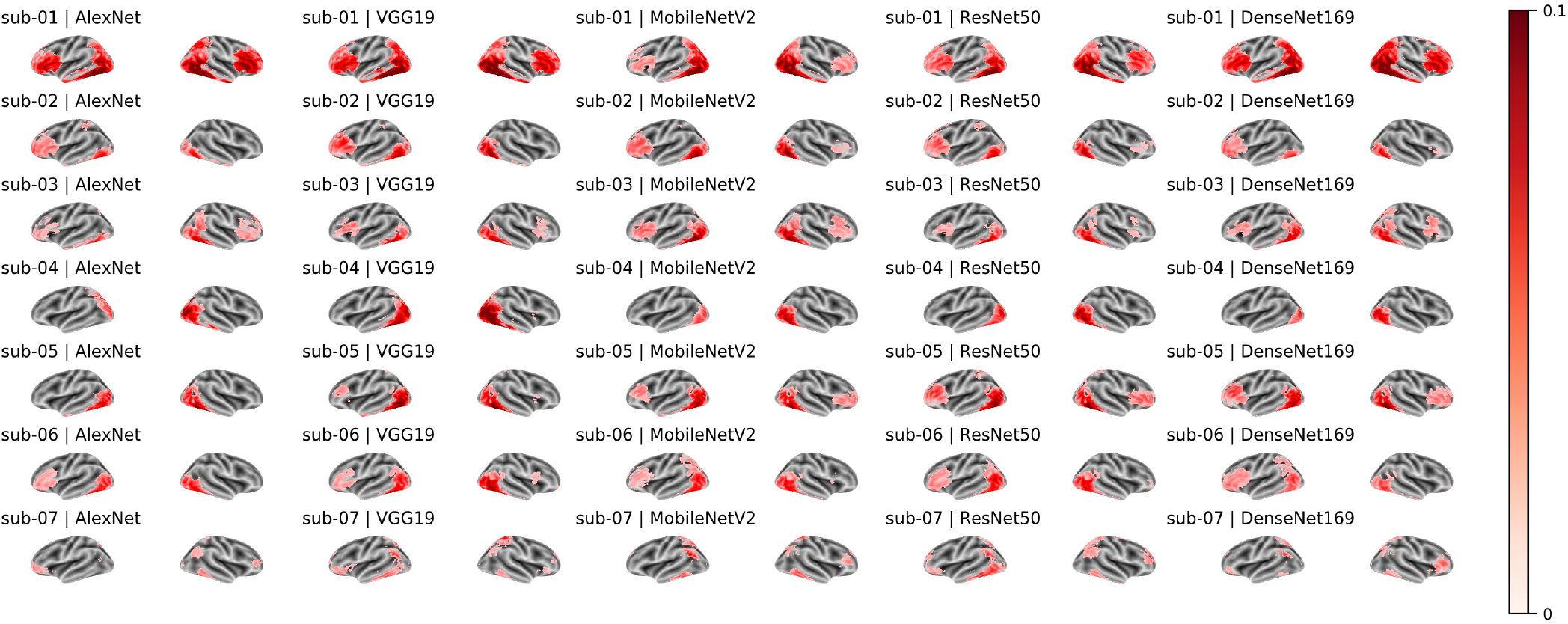
RSA map of conscious condition for the five FCNNs. Each row showed the left and right hemisphere RSA maps of the same subject, and every two columns showed the left and right hemisphere RSA maps of the same computer vision model.

### Encoding-based searchlight RSA results

Compared to the standard whole-brain searchlight RSA, the encoding-based approach first contextualizes the model representations in the source brain state (i.e., conscious) and then measures the model-relevant representations in the target brain state (i.e., unconscious). Thus, the encoding-based approach allows us to measure the model-relevant representations with one extra dimension and test the representational similarity across brain states.

Across different models within the unconscious condition, the computer vision model feature representations were positively correlated to brain activity patterns in the visual areas in four out of seven subjects (column 1 of Figures 10 - 14). We also observed that the model feature representations of the AlexNet and DenseNet169 were negatively correlated to the brain activity patterns in frontal areas for five of seven subjects. The association between brain activity patterns and the computer vision models were much stronger and more consistent across subjects in the conscious condition. Negative correlations in RSA are difficult to interpret, thus, we only provide a descriptive and speculative interpretation in the Discussion. This pattern of results was also observed to some extent within the conscious condition (column 2 of Figures 10 - 14), with positive correlations in visual areas and negative correlations were observed beyond the visual areas. We also trained the encoding model in the conscious condition and then tested the model features to the unconscious condition (column 3 of Figures 10 - 14). Here, the pattern of results were very similar to those observed within the unconscious condition, which could be due to that the signal-to-noise ratio in the unconscious condition was very low so that the encoding models could not enhance the RSA to achieve better pattern representations.

**Figure 10:**
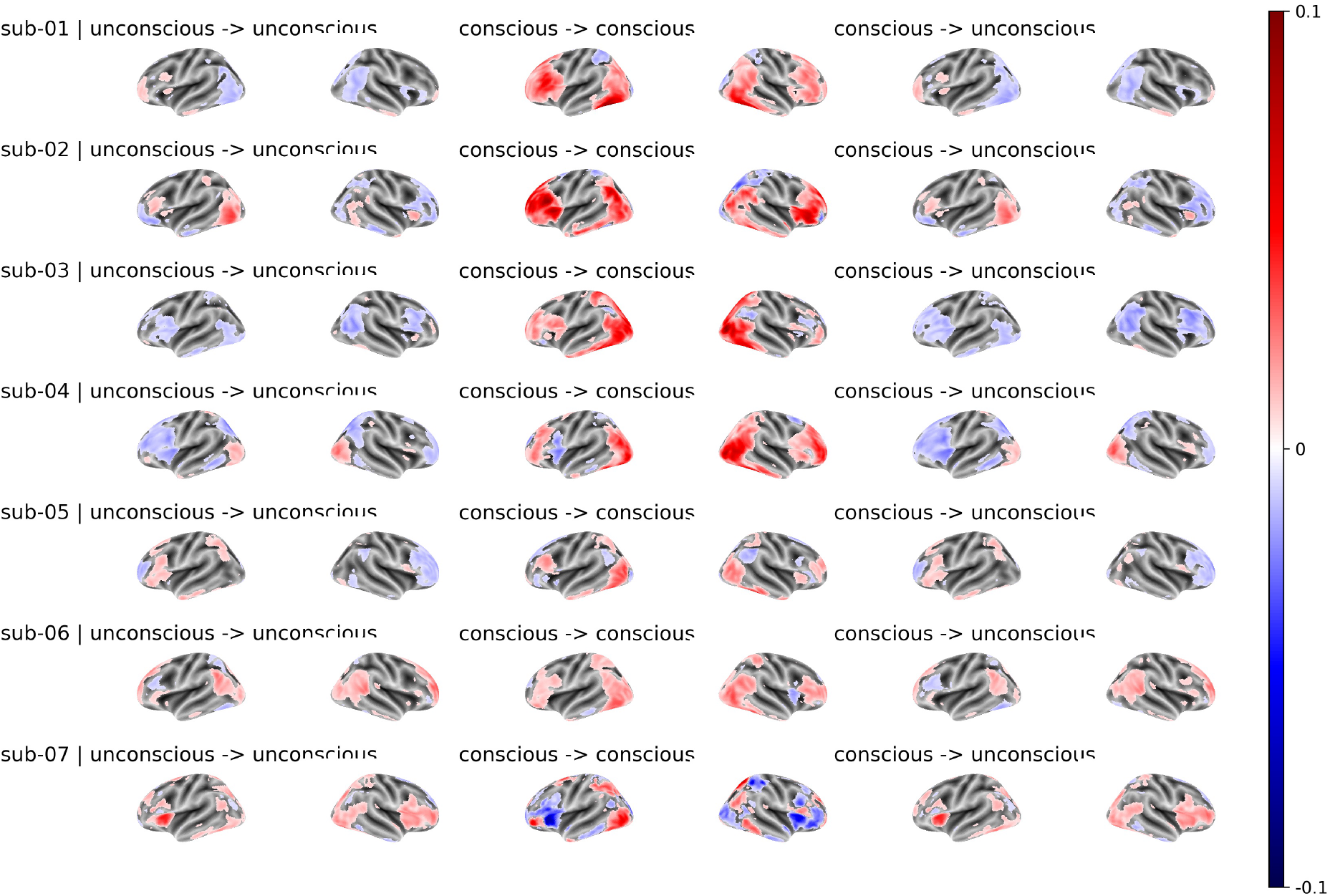
Encoding-based RSA map of the AlexNet model. The first and second columns showed the RSA maps within the unconscious condition, the third and fourth columns showed the RSA maps within the conscious condition, and the last two columns showed the RSA maps that were contextualized in the conscious condition and then generalized to the unconscious condition.

**Figure 11:**
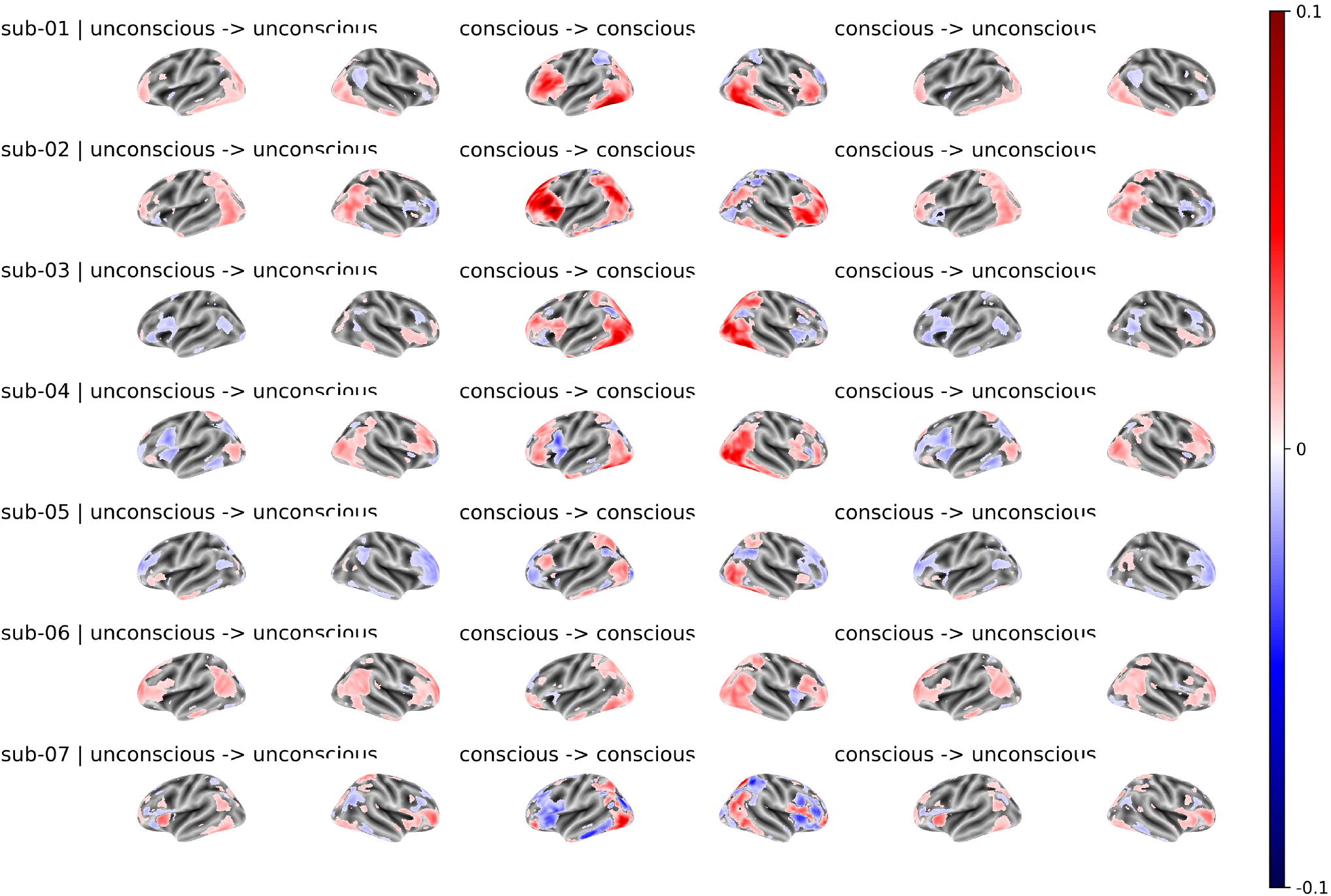
Encoding-based RSA map of the VGG19 model. The first and second columns showed the RSA maps within the unconscious condition, the third and fourth columns showed the RSA maps within the conscious condition, and the last two columns showed the RSA maps that were contextualized in the conscious condition and then generalized to the unconscious condition.

**Figure 12:**
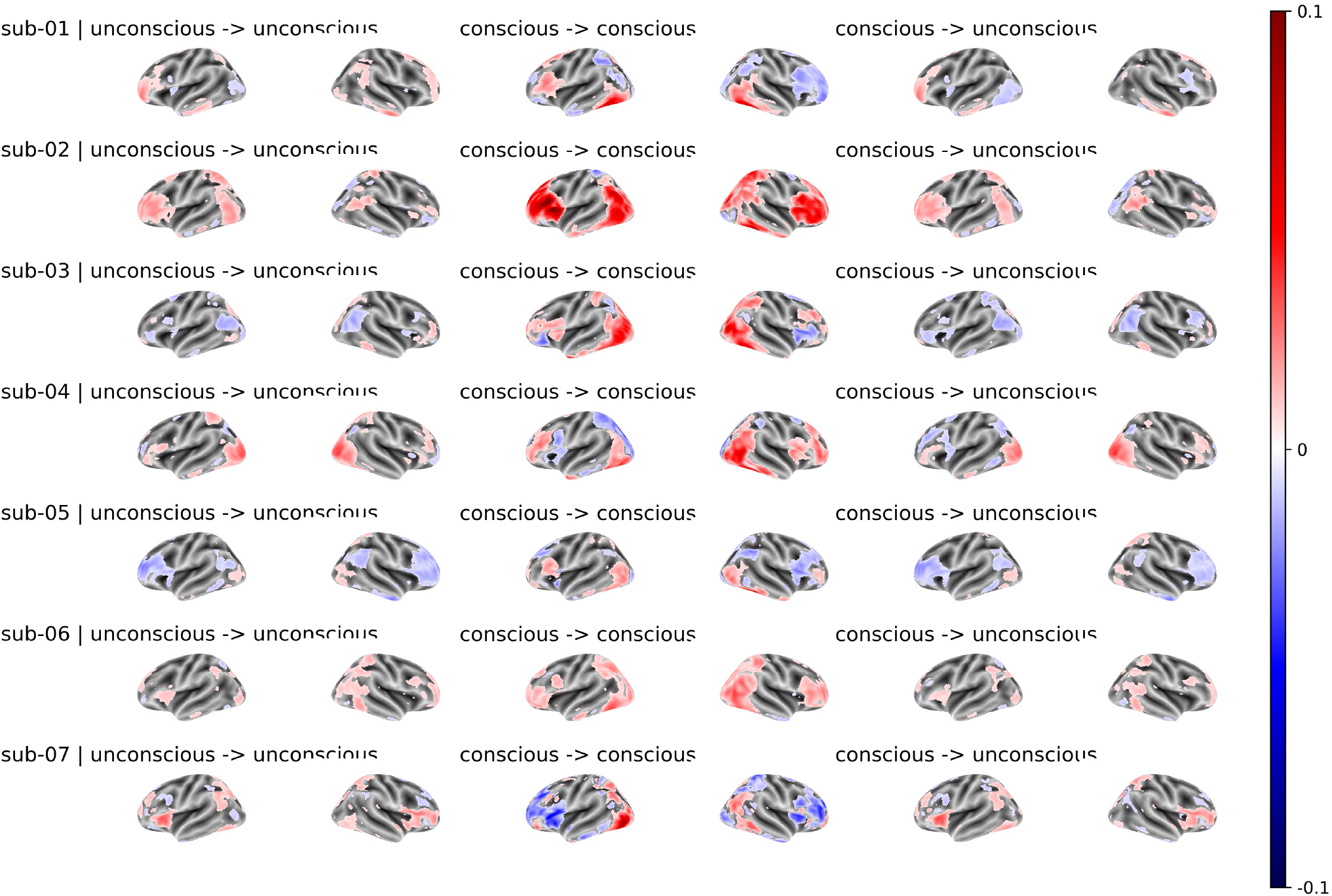
Encoding-based RSA map of the MobileNetV2 model. The first and second columns showed the RSA maps within the unconscious condition, the third and fourth columns showed the RSA maps within the conscious condition, and the last two columns showed the RSA maps that were contextualized in the conscious condition and then generalized to the unconscious condition.

**Figure 13:**
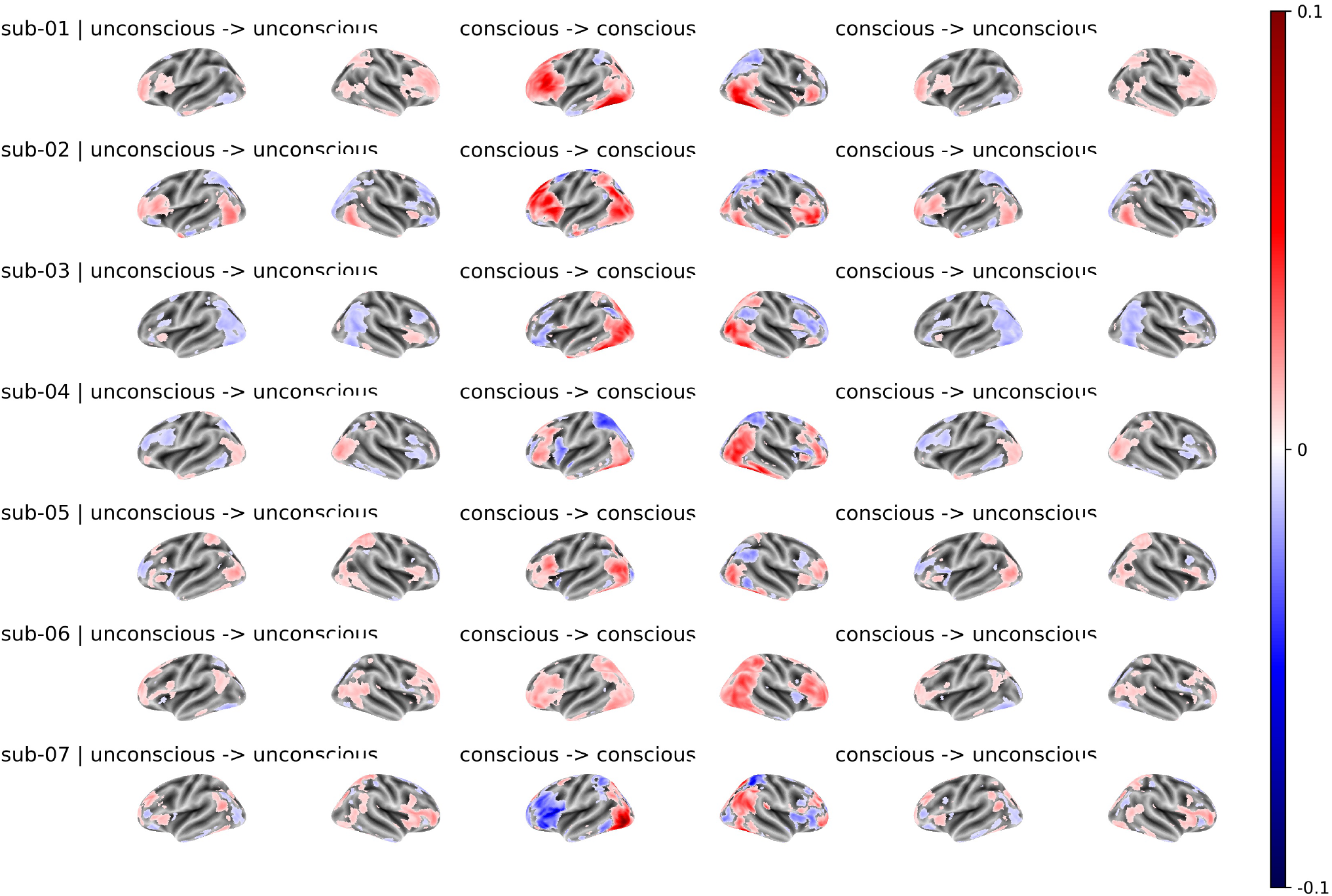
Encoding-based RSA map of the ResNet50 model. The first and second columns showed the RSA maps within the unconscious condition, the third and fourth columns showed the RSA maps within the conscious condition, and the last two columns showed the RSA maps that were contextualized in the conscious condition and then generalized to the unconscious condition.

**Figure 14:**
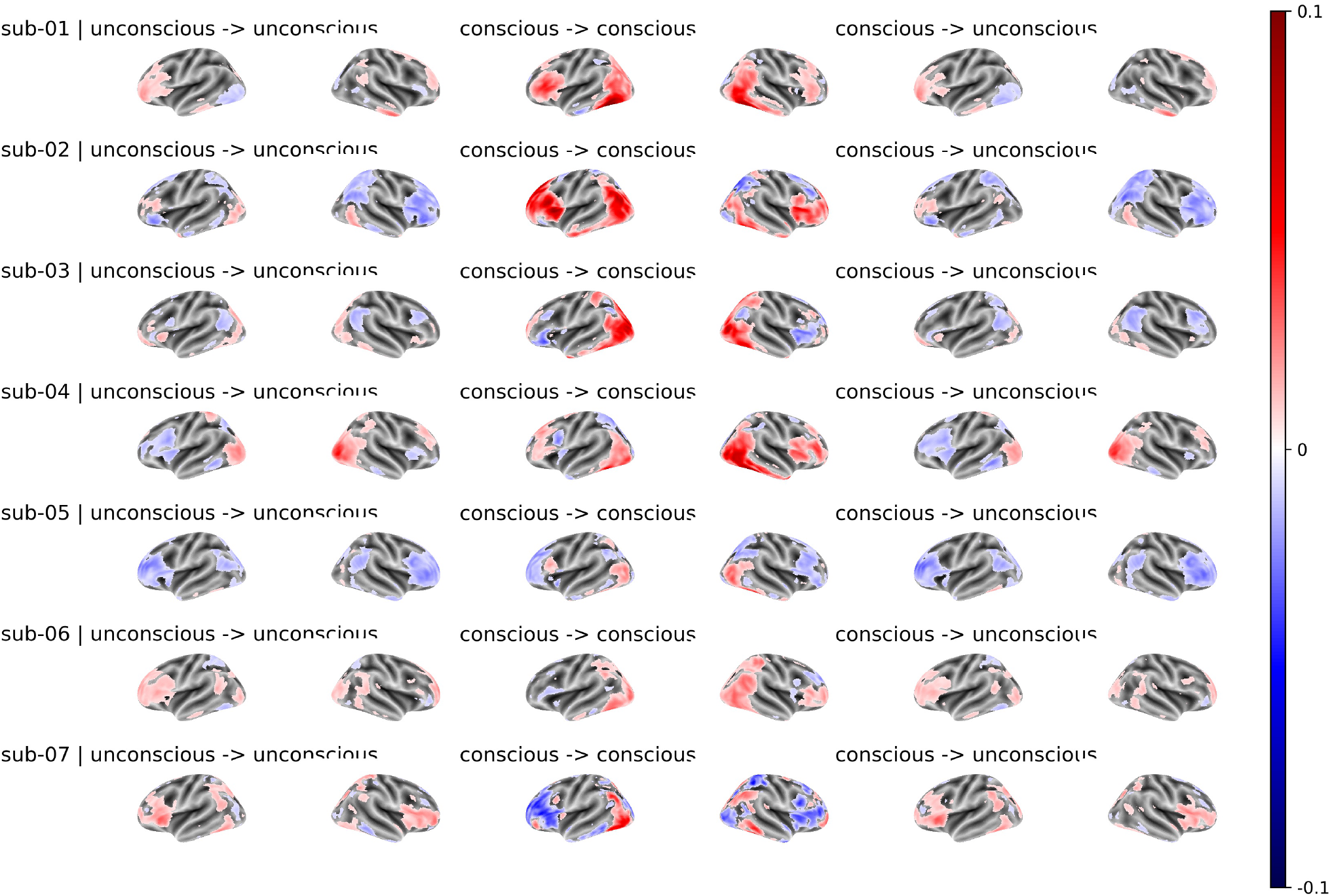
Encoding-based RSA map of the DensNet169 model. The first and second columns showed the RSA maps within the unconscious condition, the third and fourth columns showed the RSA maps within the conscious condition, and the last two columns showed the RSA maps that were contextualized in the conscious condition and then generalized to the unconscious condition.

## Discussion

We tested an information-based approach to investigate the properties of the unconscious and conscious brain representations. Specifically, we used a standard RSA pipeline ^18^ and also the encoding-based RSA ^32^ to investigate how the FCNNs’ hidden layer representations of the experimental images explained brain activity. Due to the small sample size of the subject pool and the fact that both RSA pipelines were not designed for within-subject analyses, we can only provide descriptive results from these RSA pipelines.

The standard whole-brain searchlight RSA revealed that the FCNNs’ hidden layer representations correlated to the brain representations of the unconscious contents in the ventral visual pathway in the majority of the observers, particularly for models that ranked high in explaining the variance of the visual cortex (i.e., based on brain-score, ^24^, i.e., VGGNet and ResNet50). Also five of seven subjects showed brain activity patterns that correlated with the model in frontoparietal areas in the unconscious condition. In the conscious condition, all the subjects showed neural patterns correlated with the model in visual cortex, and also in frontoparietal cortex in most subjects. Similar to the standard RSA analysis, the encoding-based RSA revealed that the brain representations of the unconscious contents in both visual and frontal areas correlated to the FCNNs’ hidden layer representations. For instance, in the VGG19 model-based RSA, six of the seven subjects showed positive correlations to the model in the visual cortex for unconscious contents. Three subjects showed positive correlations in the frontoparietal areas, and two subjects showed negative correlations to the model in the frontoparietal areas. The brain patterns correlated to the model were much stronger and more consistent across subjects in the conscious condition.

However, the results of the encoding-based RSA in the unconscious condition were mixed and some-how difficult to interpret. There were negative correlations in the visual areas for some observers in some of the models (i.e., AlexNet and DenseNet169), and there were negative correlations in the frontal areas for most of the subjects in many of the models (i.e., ResNet50 and DenseNet169). We interpret the negative correlations as showing that the brain representations were associated with the model representation but each having a different structure. While the hidden representations of the FCNNs of the images are known to explain a large variance of the brain responses ^24^, the brain mechanisms for processing information across different levels of image noise may still be quite different from the FCNNs. The FCNNs, particularly those trained with supervised methods for image classification ^35^, learn to represent the images through different layers in a bottom-up fashion, but the brain represents the images not only through bottom-up processes in the visual stream but also through top-down processes, incorporating feed-back information from other higher-order areas (i.e., prefrontal cortex). Thus, representations in the brain could be more visual-oriented or more semantic-oriented in terms of different levels of processing ^36^, which are likely different (e.g. deeper) in the human brain compared to the FCNNs. The representations in the ventral visual pathway are primarily visual-oriented like the computer vision model, thus, the pattern representations were positively correlated to the FCNNs’ hidden representations. On the other hand, when the information from visual areas is projected to higher-order areas in the frontal cortex, the representations might become more semantic-oriented in the brain but comparatively less so in the model. Hence, it is possible that the brain representations in the frontal cortex were were similar to the computer vision models, but the projected representations from the visual areas “pointed to” a different high-dimensional direction resulting in negative patter correlations in the encoding-based RSA.

## Author contributions

N.M. and D.S. designed the study; N.M. analysed the data under the guidance of R.S. and D.S. N.M. prepared a first draft of the paper. All authors discussed the results and contributed towards the writing of the final version of the manuscript; D.S. supervised the project.

## Data availability statement

Analysis scripts are available at https://github.com/nmningmei/unconfeats. The fMRI data can be found at https://openneuro.org/datasets/ds003927

## Acknowledgements

D.S. acknowledges support from the Basque Government through the BERC 2018-2021 program, from the Spanish Ministry of Economy and Competitiveness, through the ‘Severo Ochoa’ Programme for Centres/Units of Excellence in R & D (CEX2020-001010-S) and also from project grants PSI2016-76443-P and PID2019-105494GB-I00 from MINECO. R.S. acknowledges support by the Basque Government (IT1244-19 and ELKARTEK programs), and the Spanish Ministry of Economy and Competitiveness MINECO (project TIN2016-78365-R).

## Footnotes

a torch.nn.AdaptiveAvgPool2d((1,1))

b torch.nn.SELU(inplace=False)

c Scipy.spatial.distance.pdist

d sklearn.preprocessing.MinMaxScaler

e sklearn.preprocessing.StandardScaler

